# Anti-proliferative and Apoptosis Inducing Effect of Thymoquinone in Human Teratocarcinomal (NTERA-2) Cancer Stem-Like Cells

**DOI:** 10.1101/2025.09.08.674795

**Authors:** Rajitha K. Rathnayaka, Fathima T. Muhinudeen, Nirwani N. Seneviratne, Dipun N. Perera, Shalini K. Wijerathne, Umapriyatharshini Rajagopalan, Kanishka Senathilake, Dinara S. Gunasekara, Kamani H. Tennekoon, Sameera R. Samarakoon

**Affiliations:** Institute of Biochemistry, Molecular Biology and Biotechnology, University of Colombo, No. 90, Cumarathunga Munidasa Mawatha, Colombo, 00300, Sri Lanka; New Frontier Bio, 48 Danham Street, Beverly, MA 01915, USA

**Author notes:** **Corresponding author:** Sameera R. Samarakoon, Institute of Biochemistry, Molecular Biology and Biotechnology, University of Colombo, 90, Cumaratunga Munidasa Mawatha, Colombo 00300, Sri Lanka., (Ext- 318).

**Keywords:** *Nigella sativa*, cytotoxicity, apoptosis, Thymoquinone, NTERA-2

## Abstract

Cancer stem cells (CSCs) are key drivers of tumor progression, therapeutic resistance and recurrence. *Nigella sativa*, a medicinal plant widely used in traditional medicine, has gained significant importance due to its diverse pharmacological properties. Thymoquinone (TQ), a biologically known active compound isolated from *N.sativa*, has demonstrated anticancer properties in various cancers. However, its effect on CSC-like cells has not been fully elucidated. In the present study, the anti-proliferative and apoptosis inducing properties of TQ was evaluated on human embryonal carcinoma cells (NTERA-2, cancer stem cell like model) and human peripheral blood mononuclear cells (PBMCs) *in vitro*. Antiproliferative effects of TQ on NTERA-2 cells and PBMCs were evaluated using the Sulforhodamine B (SRB) and WST-1 assays, respectively. The effect of TQ was further evaluated using colony formation assay, cell migration assay, fluorescence microscopy and quantification of caspase 3/7 activities. Oxidative stress markers (reactive oxygen species [ROS]) were also determined in NTERA-2 cells treated with TQ. Thymoquinone revealed promising dose- and time-dependent antiproliferative effects (half-maximal inhibitory concentration [IC_50_] 1.282, 1.167, and 0.984 *μ*g/mL at 24, 48, and 72 h post-treatment) in NTERA-2 cells while exerting a minimal cytotoxic effect in PBMCs. Apoptosis related morphological changes, and increased Caspase 3/7 activities confirmed the pro-apoptotic effects of TQ. Further, NTERA-2 cells treated with TQ expressed a significant increase (*P* < 0.001) in intracellular ROS activity. Overall results confirm that TQ exerts anti-proliferative and apoptotic effects in a dose- and time-dependent manner. Therefore, TQ can be considered as a potent drug lead for chemotherapy and radiotherapy resistant cancer stem cells.

## Introduction

*Nigella sativa*, an annual herbaceous plant, belonging to Ranunculaceae family grows in southwest Asia, Europe, North Africa and Turkey regions.^[1]^ The seeds of *N. Sativa* have been used for centuries in traditional medicine systems worldwide to treat various diseases, including cancer. Up to date, many studies have shown that *N. Sativa* and its main active ingredient, Thymoquinone (TQ), is pharmacologically effective against a variety of ailments, including many chronic diseases and cancers.^[2]^

Cancer stem cells (CSCs), a subset of cancer cells, are known to be responsible for cancer initiation, therapeutic resistance, metastasis and recurrence.^[3-4]^ Uncontrolled cell proliferation, dysregulated growth factor signaling pathways and resistance to conventional therapies make it challenging to find a permanent cure for cancer.^[5]^ Dysregulations of the signaling pathways such as Wnt/ β-catenin, Notch and Hedgehog are more common in CSCs and these aberrations highlight the urgent need to investigate new therapies that target CSCs.^[6]^

It has been reported from many previous studies that TQ demonstrates antitumor properties against many cell types such as human colorectal, myeloblastic leukemia, prostate, pancreatic, hepatocellular and breast cancers *in-vitro*.^[7]^ Furthermore, TQ has demonstrated antitumor activity through the modulation of multiple molecular mechanisms^[8-9]^; induction of caspase 3/7 activities^[7]^, inhibition of cell migration and invasiveness, induction of ROS and mitochondria-mediated apoptosis^[10]^, cell cycle arrest^[11]^ and various other signaling pathways.^[12]^

Anti-proliferative and apoptotic effects of TQ on NTERA-2 cl.D1 teratocarcinomal (NTERA-2) cancer stem cell like cells, have not yet been evaluated so far. NTERA-2 cell line is a completely characterized highly pluripotent cancer stem cell line^[13]^, and it has been recognized as the most convenient model to investigate the fundamental molecular mechanisms of embryonic stem cells or CSCs.^[14]^ The present study was designed to investigate the potential, anti-proliferative and pro-apoptotic effects of TQ in NTERA-2 cells, a cancer stem cell model.

## Materials and Methods

### General

All chemicals used in this study were purchased from Sigma-Aldrich (St. Louis, MO, USA) unless otherwise specified. Human embryonal carcinoma (NTERA-2 cl.D1; CRL-1973) cell line and reagents used for cell culture were purchased from American Type Culture Collection (ATCC; Manassas, VA, USA). Caspase 3/7^®^ assay kit was purchased from Promega (Promega Corporation, Madison, WI, USA). The WST-1 cell proliferation assay kit was purchased from Abcam (Cambridge, MA, USA).

### Cell culture

NTERA-2 cells were cultured in Dulbecco’s Modified Eagle Medium (DMEM), according to the ATCC recommendations, supplemented with 10% fetal bovine serum (FBS), 50 IU/mL penicillin, and 50 μg/mL streptomycin in 25 cm^2^ cell culture flasks. Cells were incubated at 37°C in 5% CO_2_ and 95% humidity.

### Suforhodamine B (SRB) assay

Suforhodamine B (SRB) assay was performed to evaluate the anti-proliferative effect of TQ in NTERA-2 cells according to the method previously described by Samarakoon et al.^[15]^ with slight modifications. Upon 80% confluence, cells were trypsinized and seeded into 96-well plates (5000 cells/well) and incubated for 24 h. After 24 h incubation, the cells were exposed to different concentrations of TQ (1.5625, 3.125, 6.25, 12.5, 25.0 µg/mL) for 24 h, 48 h and 72 h incubation periods. Paclitaxel (0.0625, 0.125, 0.25, 0.5, 1 µg/mL) was used as the positive control. After the incubation periods, cells were washed three times with phosphate buffered saline (PBS) and fixed with ice-cold 50% Trichloroacetic Acid (TCA). Plates were incubated at 4 °C for 1 h and washed five times with tap water. Fixed cells were stained with 0.4% SRB dye for 20 min at room temperature in the dark. Unbound dye was removed by washing the cells with 1% acetic acid. The protein bound dye was solubilized by adding 100 μL of unbuffered tris base solution and the plates were shaken for 1 h at room temperature. The absorbance readings were obtained using a microplate reader (Synergy HT microplate reader, BioTek Instruments, USA) at 540 nm wavelength. IC_50_ was calculated using Graph Pad Prism 7.0 (GraphPad Software Inc., San Diego, CA, USA). Untreated control contained 0.1% DMSO and ATCC recommended cell culture medium (Dulbecco’s Modified Eagle Medium [DMEM]).

### Determination of cell-morphological changes related to apoptosis

NTERA-2 cells (2 × 10^5^ cells/well) were seeded in 96-well plates and incubated for 24 h at 37°C in 5% CO_2_. The cells were then exposed to five different concentrations of TQ ranging from 1.5625 - 25 µg/mL for 24, 48 and 72 h at 37°C in 5% CO_2_. 0.1% DMSO in DMEM medium was used as the untreated control. Morphological changes were observed using an inverted phase contrast microscope (Olympus CKX41SF, Japan) post incubation.^[16]^

### Isolation of Peripheral Blood Mononuclear Cells and evaluation of antiproliferative effect by using WST-1 assay

Peripheral blood mononuclear cells (PBMC) were isolated according to the method described by Tharmarajah et al.^[17]^, and was used as the non-tumorigenic control cells to assess the antiproliferative effect of TQ and Paclitaxel. Human blood was collected and diluted using Hank’s Buffered Salt Solution (HBSS) and layered upon Histopaque-1077. Diluted and layered blood was centrifuged at 400 g for 30 min at room temperature. Afterwards the opaque interface containing mononuclear cells was carefully transferred to a clean conical centrifuge tube and cells were washed with 10 mL of isotonic PBS. Roswell Park Memorial Institute 1640 (RPMI) medium supplemented with 10% FBS, 50 IU/mL penicillin and 50 µg/mL streptomycin was used to resuspend the PBMCs. Cells were seeded in 96-well plates (5 × 10^4^ cells/ well) and incubated for 24 h. The cells were treated with five different concentrations of TQ (1.5625 - 25.0 µg/mL) and Paclitaxel (0.0625 - 1 µg/mL) for 24 h, 48 h and 72 h. The water-soluble tetrazolium (WST)-1 cell proliferation assay was performed according to the manufacturer’s instructions to determine the antiproliferative effects. DMSO (0.1%) in RPMI medium was used as the untreated control.

### Colony Formation Assay

The assay was performed according to the method described by Franken et al.^[18]^ with further modifications described by Abeysinghe et al.^[19]^ NTERA-2 cells (500 cells/mL) were seeded in 96-well plates and incubated for 48 h. Cells were treated with different concentrations of TQ (1.5625 - 25.0 µg/mL) and Paclitaxel (0.0625 - 1 µg/mL) and incubated for 7 days. Post-incubation period, the cell colonies were counted after SRB dye staining. Colony formation efficiency was calculated using the following equation: ([Number of colonies / No. of seeded cells] x 100%).

### Cell Migration Assay (Wound healing assay)

The assay was performed to evaluate the effect of TQ and Paclitaxel (positive control) on the migration rate of NTERA-2 cells. Cells (2 × 10^5^/ well) were seeded in 24-well plates and incubated up to confluence. A vertical wound was made through the cell monolayer using 200 µL pipette tips under sterile conditions as described by Wang et al.^[20]^ The cell debris was removed by washing the wells using cell culture medium. A concentration series of TQ (0.5, 1, 2 μg/mL) and Paclitaxel (0.25, 0.5, 1 μg/mL) were selected, based on the 48 h IC_50_ value, and added to each well. Treated cell culture plates were incubated at 37°C in 95% air and 5% CO_2_ atmosphere under humidified conditions. The wound was observed at different time intervals using an inverted phase contrast light microscope. Morphological observations of each well before treatment (initial), just after treatment and at different time intervals were observed for wound closure and pictures were taken. The width of the cell free area in the confluent monolayer was measured using the scale bar to quantify the ability of TQ to modulate cell migration rate.

### Fluorescence microscopic analysis

NTERA-2 cells (2 × 10^5^ cells/mL) were cultured on cell culture-treated coverslips and incubated for 24 h. Cells were then exposed to different concentrations of TQ (0.25, 0.5, 1, 2 µg/mL), dissolved in DMSO (0.1%) and incubated for 48 h. DMSO (0.1%) in DMEM medium was used as the untreated control. Incubated cells were fixed using 4% formaldehyde in the dark for 10 min. Thereafter, the cells were stained with acridine orange (AO), ethidium bromide (EB) and Hoechst 33258 dye. Excess dye was removed by washing with PBS and the cell images were captured using a fluorescence microscope (Olympus BX 51 TRF, Japan) to observe apoptotic features as described by Ediriweera et al.^[21]^

### Caspase Glo® 3/7 assay

NTERA-2 cells (2× 10^4^ cells/well) were seeded in 96-well plates and incubated for 24 h. Then the cells of each well were treated with different concentrations of TQ and Paclitaxel (in 0.1% DMSO) (0.25, 0.5, 1 *μ*g/mL) and further incubated for 48 h. The concentrations of TQ and paclitaxel used for treating was kept below the 48 h IC_50_ values. 0.1% DMSO in DMEM was used as untreated control for the experiment. Caspase3/7 reagent (100 μL) was added to each well and incubated for 1 h in the dark at 37°C. Caspase Glo**®** 3/7 (Promega) assay was performed according to the manufacturer’s instructions to evaluate caspase activity.

### Nitroblue tetrazolium (NBT) reactive oxygen species (ROS) assay

This assay was carried out on NTERA-2 cells treated with TQ according to previously described methods^[22]^ with slight modifications. NTERA-2 cells (20,000 cells/well) were cultured in 96 well plates and incubated for 24 h. Thereafter, the cells were exposed to TQ (0.25, 0.5, 1 µg/mL) and further incubated for 48 h. The concentrations of TQ used was kept below 48 h IC_50_ value. Treated cells were then incubated with 20 µL of NBT solution (1mg/mL) for 1 h at 37°C in the dark. The blue formazan precipitate was then solubilized by adding 100 µL of DMSO and the absorbance was read at 560 nm using a microplate reader.

### Statistical Analysis

All experiments were carried out in triplicates. Graph Pad Prism 8.00 (GraphPad Software Inc., San Diego, CA, USA) was used to perform the statistical analysis and results were expressed as mean ± standard deviation. Half-maximal inhibitory concentrations (IC_50_) were obtained through linear regression analysis. One-Way Analysis of Variance (ANOVA) with Tukey’s post-hoc test was used to compare groups and *p* < 0.05 was considered to be statistically significant. One-way analysis of variance (ANOVA) with Bonferroni post-hoc test was used to evaluate the effect of TQ and Paclitaxel on Caspase 3/7 expression.

## Results

### Anti-proliferative effects of Thymoquinone

#### Effect of Thymoquinone on cell proliferation in NTERA-2 and peripheral blood mononuclear cells (PBMC)

The antiproliferative activity of TQ was tested against NTERA-2 cells and PBMC at concentrations of 1.5625, 3.125, 6.25, 12.5, and 25 µg/mL, for a period of up to 72 h. Paclitaxel was used as the positive control at concentrations of 0.0625, 0.125, 0.25, 0.5, 1 µg/mL. As observed from the results of the SRB assay (Table 1), TQ exerts a potential dose and time dependent inhibition on NTERA-2 cell proliferation with IC_50_ values of 1.282, 1.167 and 0.984 µg/mL at 24, 48 and 72 h, respectively. Paclitaxel (positive control) exhibited IC_50_ values of 4.97, 1.66 and 0.592 µg/mL on NTERA-2 cells at 24, 48 and 72 h, respectively. This observation proved TQ to exert a potential anti-proliferative effect on NTERA-2 cells similar to the positive control. Further, both TQ and Paclitaxel expressed a relatively less cytotoxic effect in noncancerous peripheral blood mononuclear cells (PBMC), as indicated by the WST-1 cell proliferation assay (Table 1) with TQ exhibiting IC_50_ values of 117.3, 87.08 and 52.49 µg/mL and Paclitaxel exhibiting 230.3, 192.7 and 114.5 µg/mL at 24, 48 and 72 h, respectively.

**Table 1:** IC_50_ values (μg/mL) of thymoquinone and paclitaxel in NTERA-2 and peripheral blood mononuclear cells (PBMC)

| Cell type | Thymoquinone ($\mu\text{g/mL}$) | | | Paclitaxel ($\mu\text{g/mL}$) | | |
| --- | --- | --- | --- | --- | --- | --- |
|  | 24 h | 48 h | 72 h | 24 h | 48 h | 72 h |
| NTERA-2 cells | 1.282 | 1.167 | 0.984 | 4.97 | 1.66 | 0.592 |
| PBMC | 117.3 | 87.08 | 52.49 | 230.3 | 192.7 | 114.5 |

#### Colony Formation Assay

Thymoquinone significantly affected the colony formation ability of NTERA-2 cells in a dose dependent manner when compared to the untreated control (*P* < 0.0001) (Figure 1 (A2)). In contrast, all the concentrations of Paclitaxel (positive control) treated cells showed 0% of colony formation rates (*P* < 0.0001) (Figure1 (B2)) due to reproductive cell death. The colonies were observed under an inverted phase contrast light microscope upon staining with SRB dye (Thymoquinone treated (Figure1 (A1)) and Paclitaxel treated (Figure1 (B1)). Single cells treated with TQ lost their ability to divide and proliferate. As shown by the data, TQ can strongly inhibit the colony formation ability of NTERA-2 cells in a dose and time dependent manner compared to Paclitaxel.

**Figure 1.**
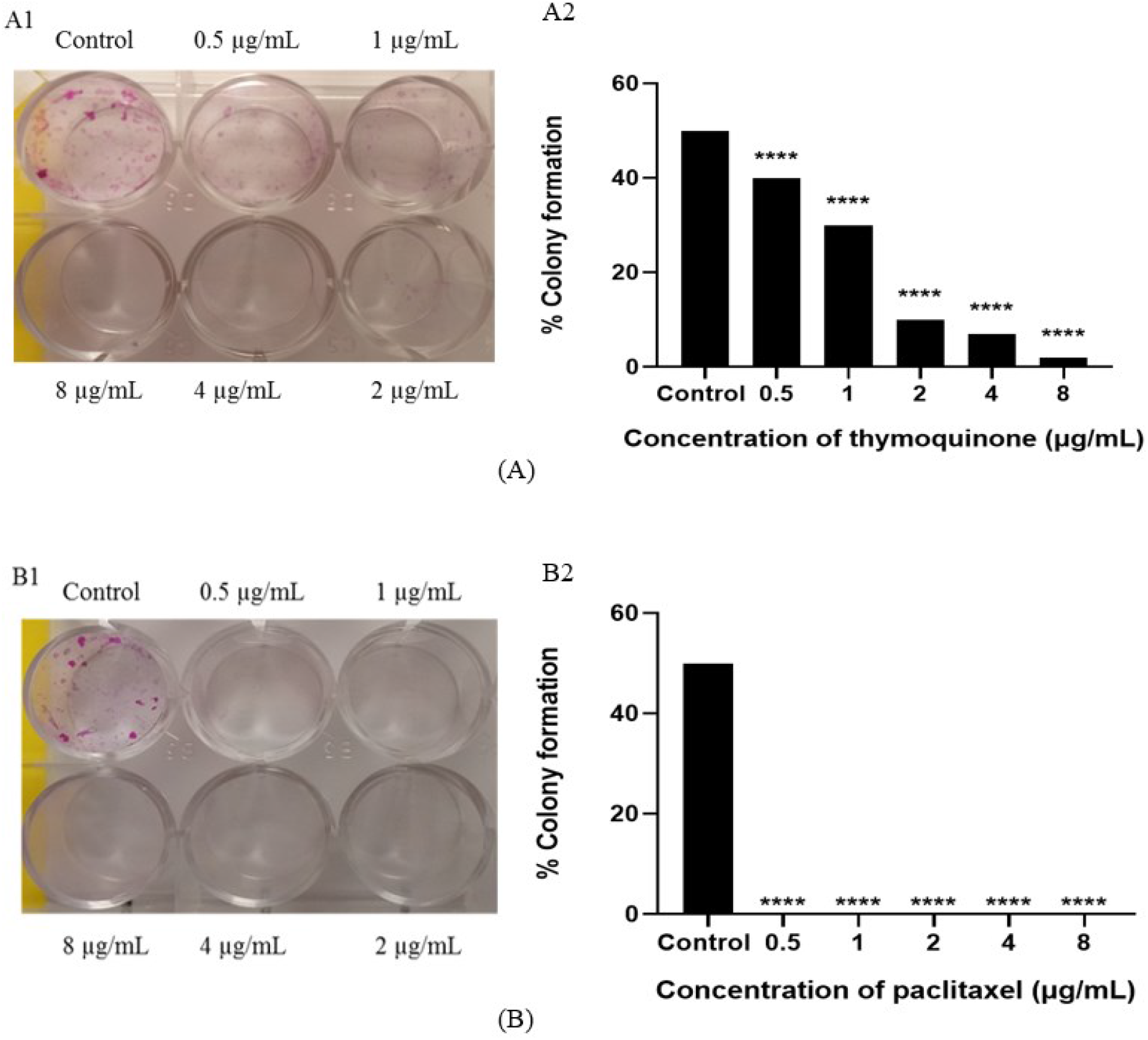
(A) Appearance of wells in a 96-well plate containing NTERA-2 cells fixed with TCA and stained with SRB dye after seven days of thymoquinone treatment (A1), with average colony formation rates (A2). (B) Appearance of wells in a 96-well plate containing NTERA-2 cells fixed with TCA and stained with SRB dye after seven days of paclitaxel treatment (B1), with average colony formation rates (B2). ****P < 0.0001.

#### Cell migration assay

Analysis of migration rate through wound healing assay revealed considerable amount of details about the migration behaviors of NTERA-2 cells treated with TQ. Dose dependent cell migration rates were observed in TQ treated NTERA-2 cells (Figure 2A) when compared to the untreated control cells. No noticeable alterations were observed at the monolayer gap created, with respect to time and dose, by Paclitaxel treated NTERA-2 cells (Figure 2B).

**Figure 2.**
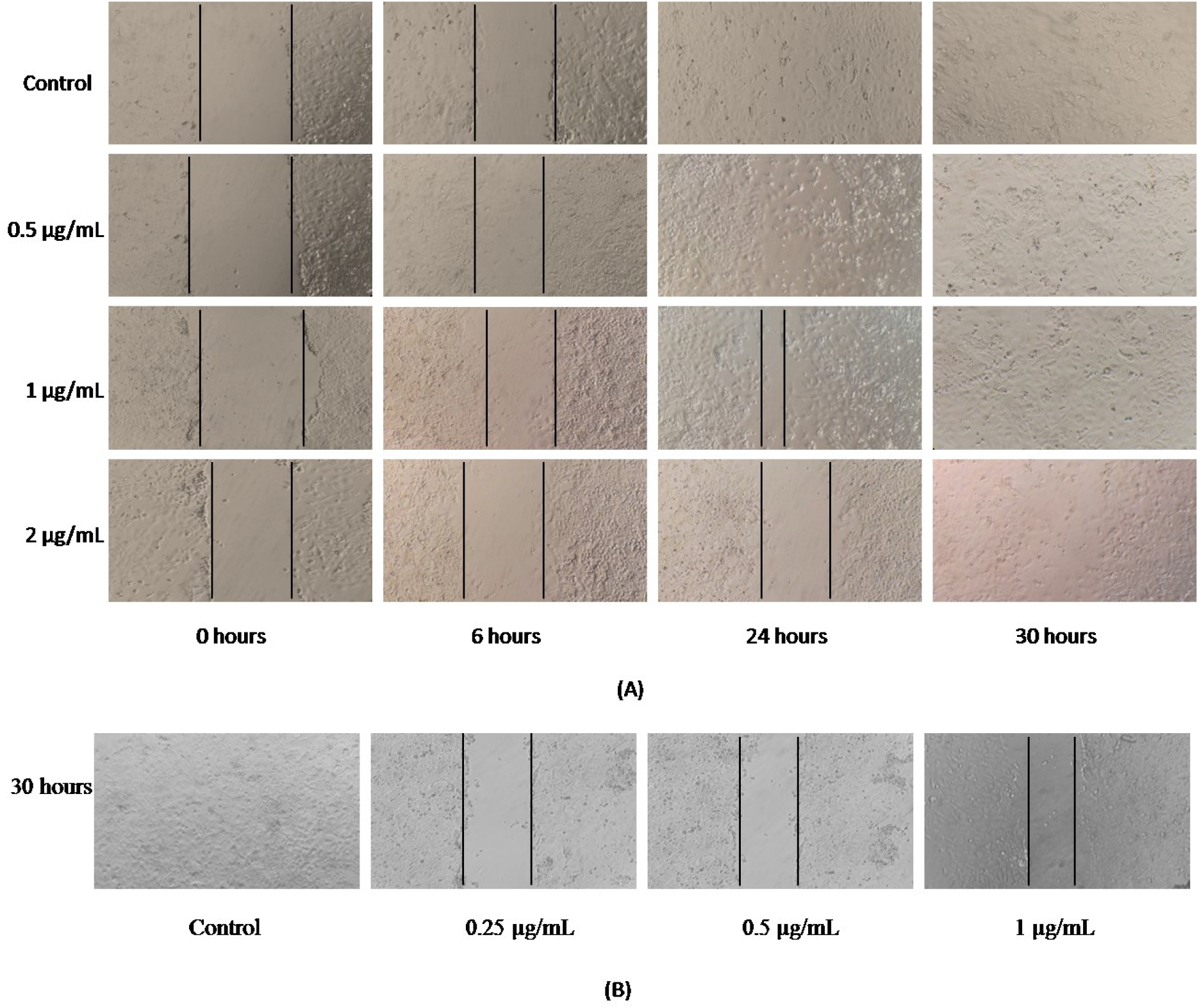
Phase contrast light microscopic images captured at different time points after creating the gap, followed by (A) thymoquinone and (B) paclitaxel treatments.

### Induction of apoptosis by Thymoquinone in NTERA-2 cells

#### Acridine orange (AO), Ethidium Bromide (EB) and Hoechst 33258 staining

Fluorescence microscopic observations of TQ treated NTERA-2 cells confirmed characteristic morphological changes related to apoptosis such as cell rounding, chromatin condensation, blebbing of plasma membrane, nuclear fragmentation, apoptotic body formation, cell shrinkage and reduction in cell volume (Figure 3). Untreated control cells exhibited a uniform green color with AO/EB staining, while apoptotic cells in the early stage showed granular yellow-green staining, and those in late stage displayed orange nuclei which were concentrated and asymmetrically localized (Figure 3a). The change in color indicates the degree of cell membrane loss 24 h post incubation with TQ in a dose-dependent manner. In the untreated control cells, uniform dispersion of Hoechst 33258 stain was observed, whereas in treated cells the stain appeared brighter and chromatin condensation was observed (Figure 3b).

**Figure 3.**
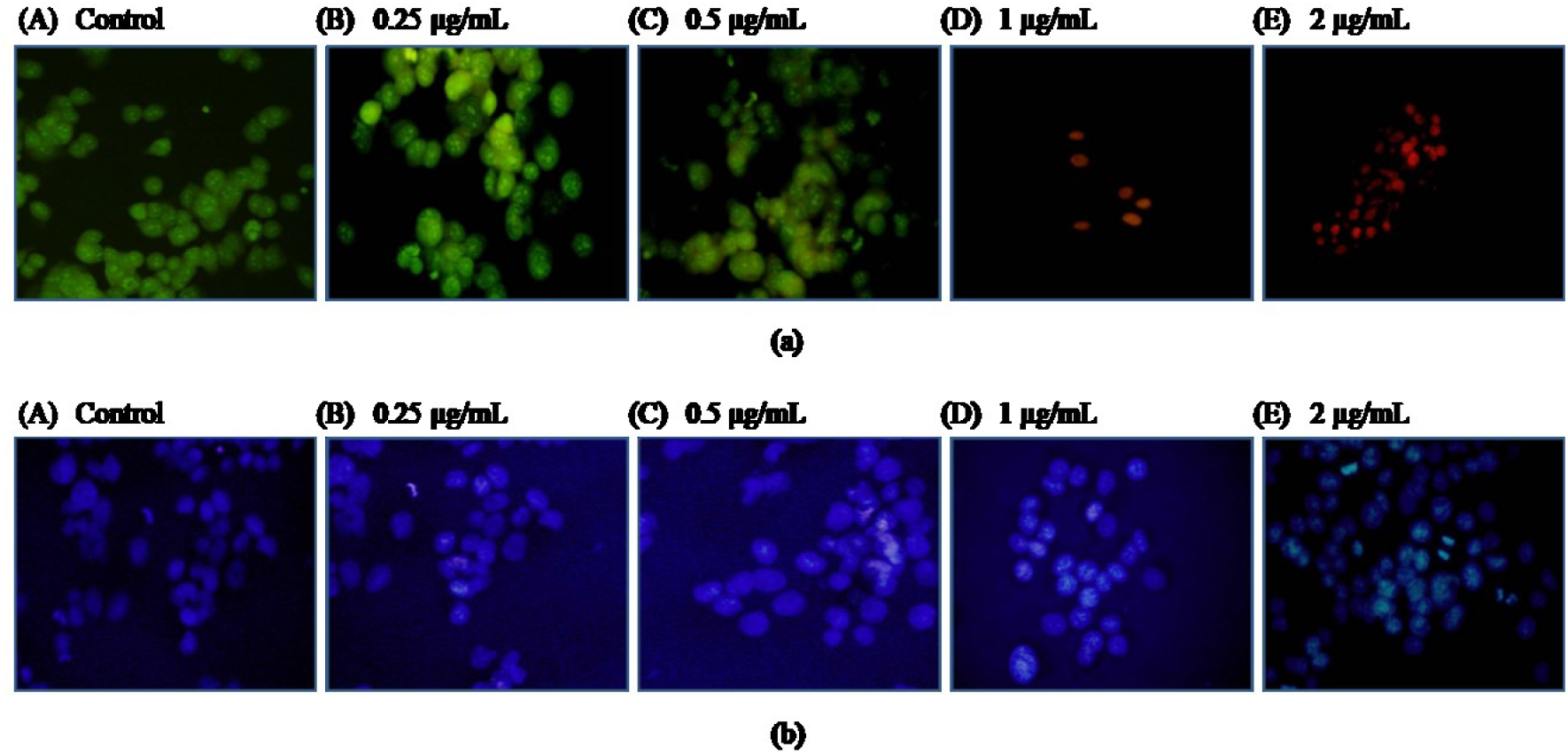
Fluorescence microscopic observations of NTERA-2 cells exposed to thymoquinone for 48 h. (a) Acridine orange (AO) and Ethidium bromide (EB) and (b) Hoechst 33258 dye (using a blue filter) (magnification 200x): A - untreated control (0.1% DMSO); B - thymoquinone (0.25 μg/mL); C - thymoquinone (0.5 μg/mL); D - thymoquinone (1 μg/mL); E - thymoquinone (2 μg/mL)

#### Effect of Thymoquinone on expression of Caspase 3/7 in NTERA-2 cells

Activities of Caspase 3 and 7 significantly (*p* < 0.0001) increased in NTERA-2 cells (Figure 4) after being exposed to TQ for 24 h. Dose dependent activation was observed at two doses (0.5 and 1 µg/mL) when compared to the untreated control.

**Figure 4.**
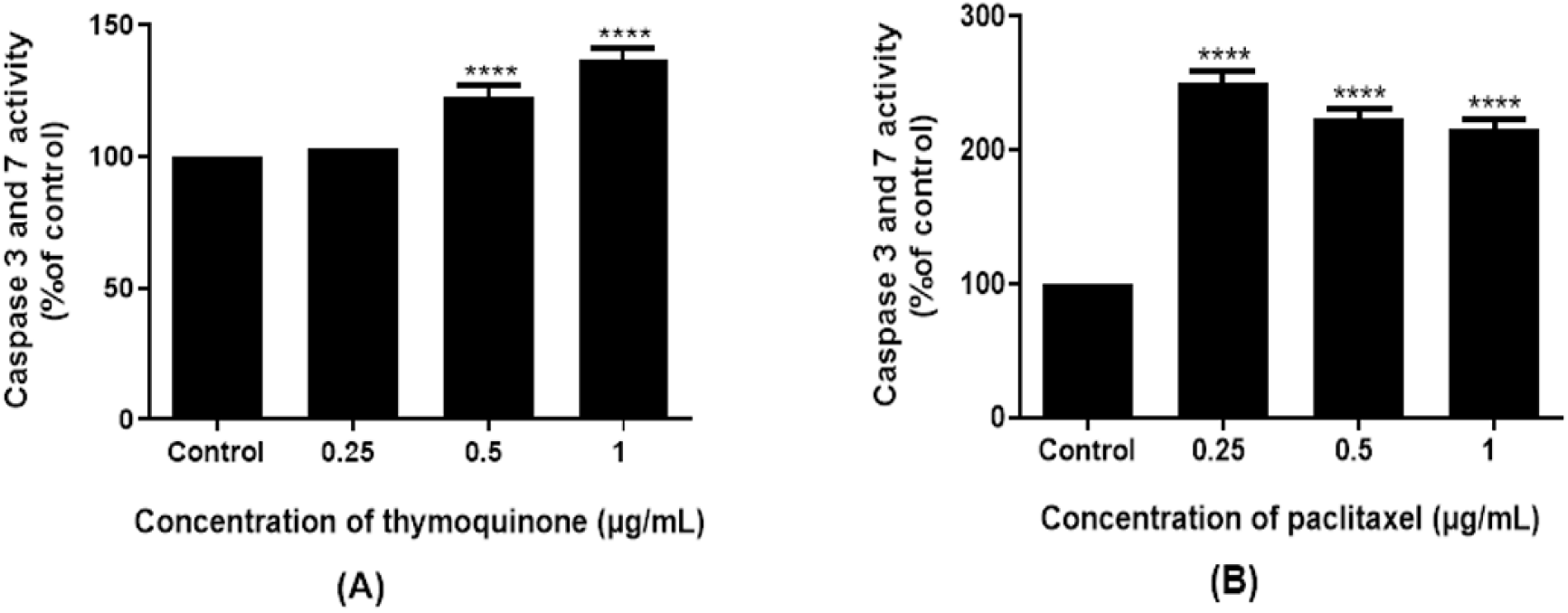
Expression of caspase 3 and caspase 7 in NTERA-2 cells after 48 hours of thymoquinone and paclitaxel (b) treatments. \*\*\*\**P* < 0.0001.

#### Effect of Thymoquinone on intracellular Reactive Oxygen Species levels

A significant (*p* < 0.0001) increase in ROS production was observed in TQ treated NTERA-2 cells at 24 h post incubation at concentrations of 0.25, 0.5 and 1 µg/mL, in comparison to the untreated control (Figure 5). The results obtained by the NBT assay demonstrate dose dependent ROS production in TQ treated NTERA-2 cells.

**Figure 5.**
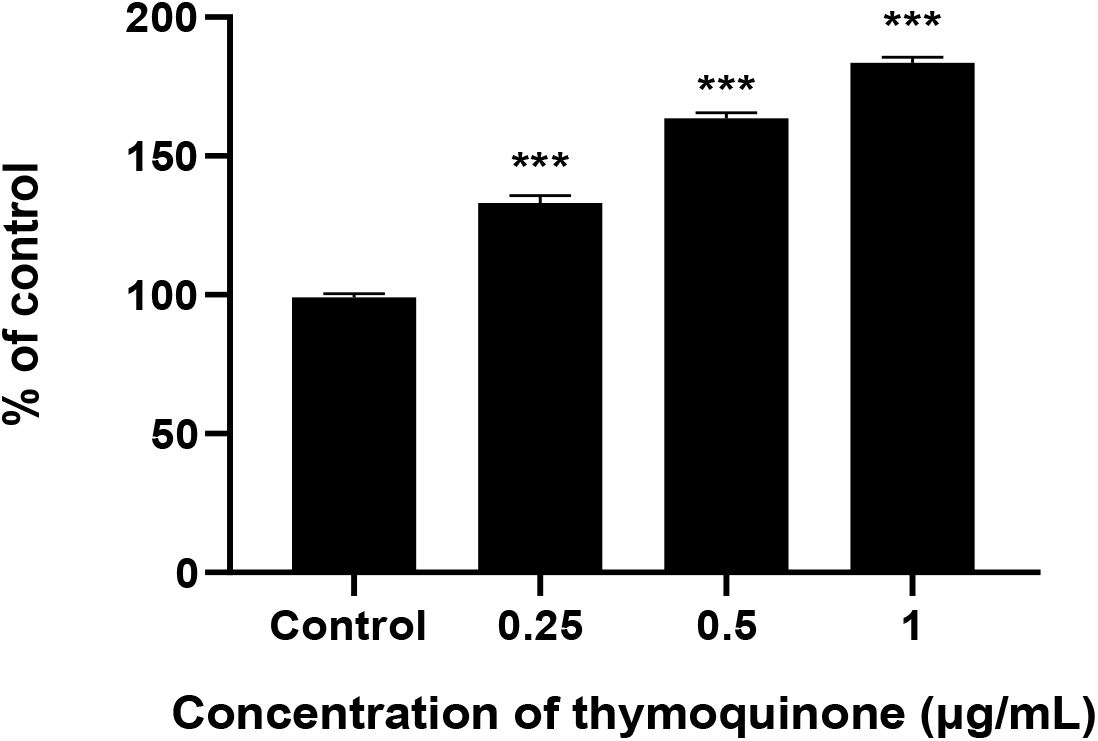
Expression of reactive oxidative stress levels after treatment of NTERA-2 cells with thymoquinone for 48 h. ***P < 0.001.

## Discussion

Thymoquinone, the major active ingredient of *N. sativa*, has been widely reported for its antioxidant, anti-inflammatory and anticancer properties.^[2]^ In this study, the anti-proliferative, apoptosis-inducing, and migration-inhibiting effects of TQ were evaluated on NTERA-2 cells (a CSC model). The experimental results from the various *in vitro* assays conducted in this study demonstrate that TQ exerts strong antiproliferative and pro-apoptotic effects on NTERA-2 cells, highlighting its potential in targeting CSCs.

A major hallmark of cancer is uncontrolled cell growth and division resulting from the dysregulation of cell cycle signaling pathways.^[23]^ Potent antiproliferative effect of TQ in NTERA-2 cells were confirmed by antiproliferative assays. In the present study, the SRB assay, used to quantitate cellular protein content^[24]^, showed a dose- and time-dependent inhibition on NTERA-2 cell proliferation when treated with TQ (Table 1). These results are consistent with previous studies, where a standardized aqueous and ethanolic extract of a polyherbal mixture containing *N. sativa* seeds exhibited strong, dose-dependent antiproliferative effects on human hepatocellular carcinoma (HepG2) cells *in vitro*. HPLC analysis confirmed the presence of TQ in the ethanolic extract of the said polyherbal mixture.^[25]^ Interestingly, a commercially available polyherbal nutraceutical capsule (Vernolac), which contains *N. sativa* seeds, demonstrated a strong dose- and time-dependent inhibitory effect on NTERA-2 cell proliferation.^[26]^ The WST-1 assay, used to measure mitochondrial dehydrogenase activity^[27]^ in healthy noncancerous cells (PBMC), indicated that TQ has no detrimental effects on non-cancerous cells similar to the widely used chemotherapeutic drug Paclitaxel (Table 1), highlighting the potential of TQ as an anticancer agent. These results correspond with previous studies where TQ was observed to induce the G1/S cell cycle arrest and suppress the expression of cyclin D1 and CDK4 in various cancer cell types including breast^[28]^, colorectal^[29]^, and prostate cancer.^[30]^ Importantly, the present study extends this antiproliferative activity on cancer stem-like cells, which are highly resistant to conventional chemotherapeutics.

Long-term proliferative capacity and self-renewal properties are defining features of CSCs.^[3]^ In the present study, we used the colony formation assay to determine the differences in reproductive viability between Paclitaxel (positive control) and TQ treated cells.^[31]^ A significant reduction in the number and size of colonies, in a dose dependent manner, post-TQ treatment (Figure 1A) indicates an effective suppression of clonogenic potential, closely in resemblance to the positive control treatment (Figure 1B). This observation is highly significant as the eradication of clonogenic CSCs is vital for the prevention of tumor recurrence.^[3]^

The cell migration assay defines the ability of cells to migrate and close the gap created artificially in a confluent monolayer.^[19]^ It was observed that TQ significantly impaired the migration rate of NTERA-2 cells. CSCs are known to contribute to metastasis through epithelial-mesenchymal transition (EMT) and enhanced motility.^[32]^ Therefore, the inhibition of migration post-TQ treatment (Figure 2A) observed in the present study indicates that TQ has the ability to modulate metastatic potential compared to Paclitaxel (Figure 2B). Previous studies support our observation, where TQ has shown to downregulate EMT markers such as N-cadherin and vimentin, while E-cadherin has been upregulated in breast and prostate cancer cells.^[33]^ This antimigratory effect may be linked to the ability of TQ to suppress MMP-2 and MMP-9 expression and interfere with cytoskeletal dynamics.^[34]^

Apoptosis, a mechanism of programmed cell death, helps maintain homeostasis of organs and tissues.^[35]^ Fluorescence microscopy using AO/EB (Figure 3a) and Hoechst 33342 (Figure 3b) staining confirmed TQ-induced apoptosis, where morphological hallmarks of apoptosis were consistent with early and late apoptotic events. These morphological changes were further substantiated at the molecular level by significant activation of Caspase 3 and 7 (Figure 4A). Caspase 3 and 7, members of cysteine proteases, are central executioners of apoptosis and play pivotal roles in coordinating the stereotypical events that occur during apoptosis.^[36-37]^ These observations align with recent findings of significant upregulation of Caspase 3 and 7 activities in NTERA-2 cells in response to Vernolac, a commercially available polyherbal nutraceutical containing *N. sativa* seeds.^[26]^

Furthermore, a significant increase in reactive oxygen species (ROS) was observed post-TQ treatment (Figure 5). ROS are reactive chemical constituents/species containing oxygen, which can harm live cells by increasing apoptosis and cell cycle arrest in cancer cells.^[38]^ DNA, proteins, and lipids are damaged when intracellular ROS levels are elevated. Thereby, triggering mitochondrial-mediated apoptosis.^[39]^ Therefore, it is likely that production of ROS after exposure to TQ causes apoptosis in NTERA-2 cells. The role ROS play as a mediator of TQ-induced cell death is well established, as studies have reported that TQ acts as a pro-oxidant at high concentrations in cancer cells, leading to oxidative stress and apoptosis.^[40]^ In the context of CSCs, where elevated antioxidant defenses are exhibited, the overexpression of TQ-induced ROS could disrupt redox homeostasis and sensitize them to cell death.^[39]^

Overall results of this study highlights the potent anti-proliferative and pro-apoptotic effects of TQ on NTERA-2 cells, with minimal cytotoxic effects in the non-cancerous peripheral blood mononuclear cells (PBMC) and proves TQ to be a promising natural compound for targeting CSCs. The inhibition of colony formation and cell migration indicates that TQ can impair tumorigenic and metastatic potential. While Caspase 3/7 and ROS upregulation supports the activation of intrinsic apoptotic pathway, further highlighting the cytotoxic mechanism of TQ and its use as a potential candidate to overcome drug resistance and heterogeneity within tumors by targeting CSC populations.

Future studies on gene and protein expression profiles related to stemness, apoptosis and signaling pathways will elucidate the precise molecular pathways through which TQ mediates potential anticarcinogenic effects in NTERA-2 cells. Moreover *in vivo* validation is vital to assess the pharmacodynamics, bioavailability, and safety profile of thymoquinone.

## Acknowledgements

The authors would like to thank the Institute of Biochemistry, Molecular Biology and Biotechnology, University of Colombo and the National Science Foundation, Sri Lanka (Grant No: RPHS/ 2016-C07) for funding.

## Authors’ Contributions

***Rajitha K. Rathnayaka*** contributed to investigation, data curation, validation, writing–original draft and writing–review and editing. ***Fathima T. Muhinudeen*** contributed to writing–original draft and writing–review and editing. ***Nirwani N. Seneviratne, Dipun N. Perera, Shalini K. Wijerathne, Umapriyatharshini Rajagopalan*** and ***Kanishka Senathilake*** contributed to writing-review and editing. ***Dinara S. Gunasekara, Kamani H. Tennekoon*** and ***Sameera R. Samarakoon*** contributed to conceptualization, project administration, methodology, funding acquisition, supervision, and writing-review and editing.

